# Apontic regulates cell proliferation and development by activating the expression of *hedgehog* and *cyclin E*

**DOI:** 10.1101/054809

**Authors:** Xian-Feng Wang, Qian Cheng, Chong-Lei Fu, Zi-Zhang Zhou, Susumu Hirose, Susumu Hirose

## Abstract

Hedgehog (Hh) signaling pathway and Cyclin E are key players in cell proliferation and development. Hyperactivation of *hh* and *cyclin E.* has been linked to several types of cancer. However, transcriptional regulation of *hh* and *cyclin E.* are not well understood. Here we show that an evolutionarily conserved transcription factor Apontic (Apt) is an activator of *hh* and *cyclin E.* in *Drosophila*. Apt directly promotes the expression of *hh* and *cyclin E.* through its binding site in the promoter regions of *hh* and *cyclin E.* during wing development. This Apt-dependent proper expression of *hh* and *cyclin E.* is required for cell proliferation and development of the wing. Apt-mediated expression of *hh* and *cyclin E.* can direct proliferation of Hh-expressing cells and simultaneous growth, patterning and differentiation of Hh-recipient cells. The discovery of the coordinated expression of Hh and principal cell-cycle regulator Cyclin E. by Apt implicates insight into the mechanism by which deregulated *hh* and *cyclin E* promotes tumor formation.

**Summary statement:** We identified a novel role for Apontic as an important common regulator of the transcription of *hedgehog* and *cyclin E.* Our study provides important insights into the mechanism of organ development.

## Introduction

Animal development requires the organ patterning and growth. How these two processes are coordinated remains unclear. The *Drosophila* wing is an excellent model to study the regulation of gene expression during the organ patterning and cell growth. The wing disc is a sac-like structure composed of disc proper (DP) cells and peripodial epithelium (PE). During larval development, wing disc DP and PE cells proliferate extensively and are patterned, finally give rise to the adult wing (Milner et al., 1984). The Hh and Cyclin E can contribute to patterning and growth of the wing disc during development (Neufeld et al., 1998;Tabata and Kornberg, 1994).

Hh pathway is one of the major signaling pathways that control animal development from *Drosophila* to humans, which has been implicated in stem cell maintenance, cell migration, axon guidance and tissue regeneration (Beachy et al., 2004;Charron et al., 2003;Clement et al., 2007;Hochman et al., 2006). In the *Drosophila* wing disc, Hh expresses in posterior (P) compartment cells and spreads into the anterior compartment where it activates target genes such as *engrailed (en), patched (ptc), collier (col), decapentaplegic (dpp) and iroquois (iro)* (Matusek et al., 2014;Nahmad and Stathopoulos, 2009;Tabata and Kornberg, 1994) to control wing patterning. Moreover, Hh is required for transient fusion between the PE and the DP sides during regeneration of wing discs (McClure and Schubiger, 2005). Therefore, the expression of x*hh* is vital in the wing disc. In the anterior (A) compartment cells, the truncated transcription repressor Ci^R^ inhibits the transcription of *hh*. However, the underlying mechanism by which the posterior cells activate *hh* transcription is still to be determined.

Cyclin E belongs to the cyclin family, which is required for cell division (Knoblich et al., 1994). Dysregulation of *cyclin E.* correlates with various tumors, including breast cancer and lung cancer (Donnellan and Chetty, 1999;Keyomarsi et al., 1994;Moroy and Geisen, 2004). Besides, deregulated Cyclin E activity causes cell lineage-specific abnormalities, such as impaired maturation due to unregulated cell proliferation (Minella et al., 2008). In *Drosophila*, Cyclin E is essential for G1-to-S phase transition in the posterior cells of eye disc (Richardson et al., 1995). It has been reported that *cyclin E* is a potential target gene of *Hh* signaling in *Drosophila*. *Hh* pathway activates *cyclin E* transcription through its unique transcription factor Ci in the posterior cells of eye disc (Duman-Scheel et al., 2002). It is known that *Hh* pathway is turned on exclusively in the A cells near A/P boundary (Strigini and Cohen, 1997;Tabata and Kornberg, 1994). However, *cyclin E* expresses throughout the wing disc (Neufeld et al., 1998). This contradiction suggests that other factors are involved in regulating the expression of *cyclin E*. Therefore, it is fruitful to investigate the regulation of *cyclin E* in wing disc and the relationship between *cyclin E* and *Hh* pathway.

Apontic (Apt) has been identified as a transcription factor involved in development of tracheae, head, heart and nervous system (Eulenberg and Schuh, 1997;Gellon et al., 1997;Liu et al., 2003;Su et al., 1999). Apt can suppress metastasis (Woodhouse et al., 2003) and is required in the nervous system for normal sensitivity to ethanol sedation (McClure and Heberlein, 2013). Moreover, Apt participates in JAK/STAT signaling pathway to limit border cells migration (Starz-Gaiano et al., 2009;Starz-Gaiano et al., 2008;Yoon et al., 2011). However, the role of Apt in wing development is unknown.

In this study, we found that both loss of and overexpression of *apt* resulted in defect wings. Further studies demonstrated that loss of *apt* attenuated the expression of *hh* and *cyclin E*, while *apt* overexpression upregulated *hh* and *cyclin E*. In addition, we found inherent Apt binding sites in the promoter region of *hh* and *cyclin E*. Mutating the sites inhibited the expression of *hh* and *cyclin E.* Collectively, Apt activates the expression of *hh* and *cyclin E*to allow proper wing development.

## RESULTS

### Apt is expressed in the wing disc and is required for wing development

As the first attempt to investigate the function of *apt* during wing development, we analyzed *apt* expression pattern in the wing disc by immunostaining using anti-Apt antibody. In the wing disc, Apt was detected in PE cells as revealed by co-localization with a PE marker Ubx (Fig. 1A). Apt was also detected in DP cells (Fig. 1B). These data clearly demonstrate that Apt is expressed in both the PE and DP of the wing disc, suggesting its possible role in wing development.

To analyze the role of Apt during wing development, we would examine the developing wing of homozygous *apt* null mutant. However, *apt* null homozygotes die as embryos (Eulenberg and Schuh, 1997). Therefore, we induced *apt* loss of function mutant clones in the wing disc using the *FLP/FRT* system (Theodosiou and Xu, 1998). The formation of these clones resulted in a small wing with a blistered phenotype (Fig. 1D) compared with the control wing (Fig. 1C). Furthermore, RNAi-mediated knockdown of *apt* in DP cells of the wing disc resulted in a small wing, and also reduced the width between vein 3 and vein 4 (Fig. 1E,F). Given that the space between vein 3 and vein 4 is a characteristic monitor of *Hh* pathway activity in adult wings, knockdown of *apt*-mediated narrowing the space indicates that Apt possibly regulates *Hh* pathway. RT-PCR analyses showed effective knockdown of *apt* mRNA level upon *apt-RNAi* (Fig. S1). To investigate the effect when *apt* is overexpressed, we employed *MS1096-Gal4* driver to express UAS-*apt* in both the PE and DP of the wing disc. Abnormal wings were induced by overexpression of *apt* (Fig. 1G). The wing was diminished and blistered, the pattern of veins was disrupted and extra abnormal bristles were induced in the wing margin. In addition, when UAS-*apt* was expressed by a stronger gal4 (*sd-Gal4*) in DP cells, both wings and halters were lost (Fig. 1H). Taken together, the loss-of-function and gain-of-function analyses indicate that Apt is indispensable for wing development.

**Fig. 1.**
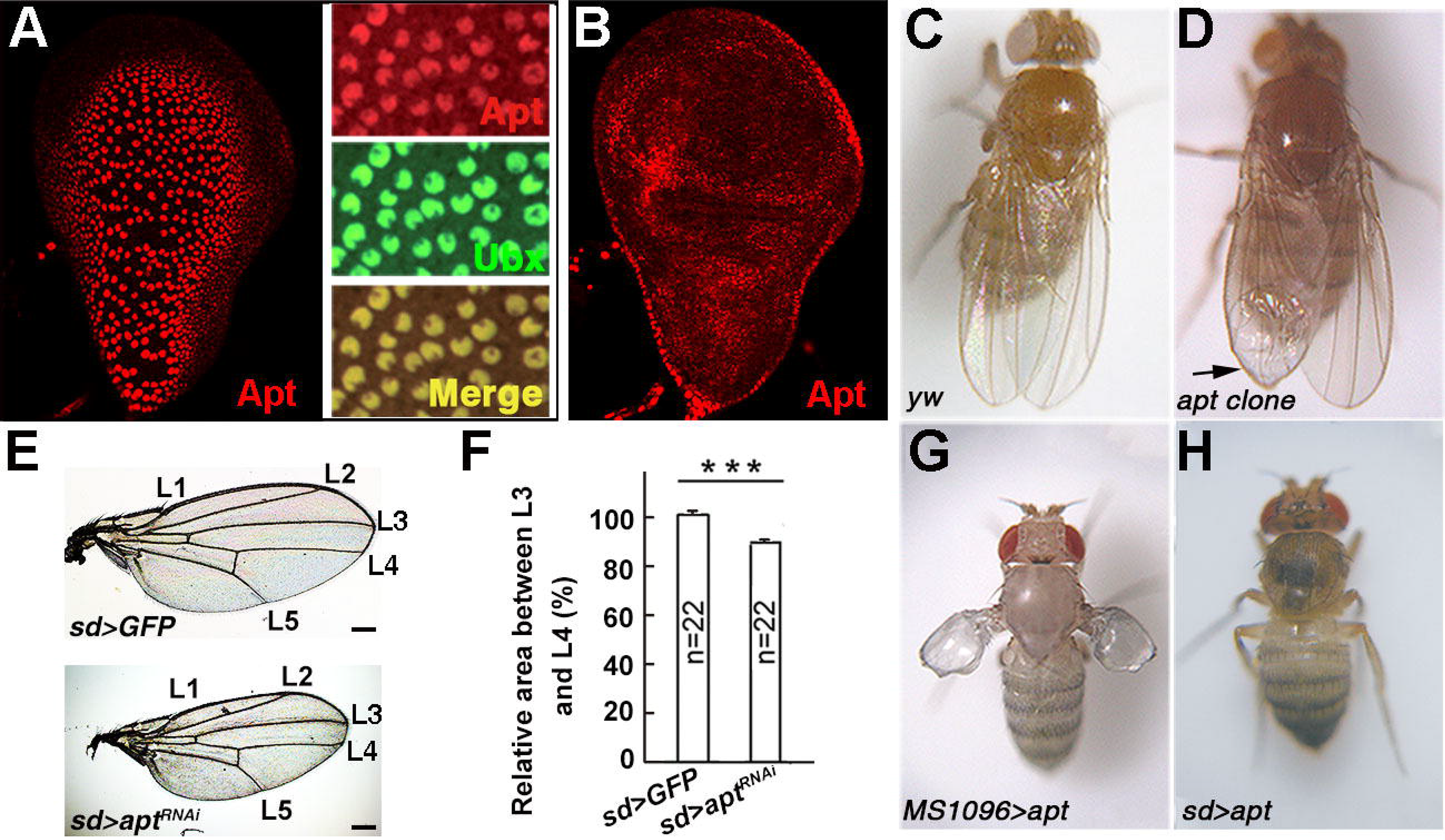
Apt is expressed in the wing disc and required for wing development.

(A) A single optical section of the PE of the wing disc from a second instar larva and the co-localization of Apt and Ubx (PE marker). (B) A single optical section of the DP of the wing disc from a second instar larva. (C) The adult wing of*yw.* (D) The adult wing of *apt^ΔA4^* clones. The arrow indicates the blistered wing. (E) The adult wings of *sd-GAL4; UAS-GFP* and *sd-GAL4*; *UAS-apt^RNAi^*. (F) Quantification of the intervein region between L3 and L4 relative to that between L1 and L2 *(sd>GFP* value was set as 100%) of *sd-GAL4; UAS-GFP* and *sd-GAL4; UAS-apt^RNAi^* by ImageJ. Error bars, SEM. Student's t tests, ***p<0.001. (G) The adult wing of *MS1096-GAL4; UAS-apt.* (H) The adult wing of *sd-GAL4; UAS-apt.* Scale bars, 200 um.

### Apt regulates the expression of *hh* in the wing disc

The observation that knockdown of *apt* narrowed the space between vein 3 and vein 4 implied that Apt might modulate *Hh* pathway in wings. As an important initiator of *Hh* pathway, *hh* gene expresses in the wing disc (Cho et al., 2000;Tabata and Kornberg, 1994).We first compared the expression of *apt* and *hh* in the wing disc, and found that Apt and *hh-lacZ* were co-expressed in PE cells (Fig. 2D-F) and posterior compartment cells of the DP (Fig. 2G-I) in the second instar larval disc. Furthermore, *apt* exhibited genetic interaction with *hh*. Transheterozygotes of two sets of *hh* alleles (*hh^bar3^/hh^2^* and *hh^Mir^/hh^2^*) exhibited smaller wing with an extra crossvein (Fig. S2), demonstrating that it is a phenotype of *hh* mutant. While wings of animals heterozygous for *hh*^2^ or *apt*-null allele were normal, transheterozygotes of *apt*-null allele and *hh*^2^ showed the same wing phenotype (Fig. 2A-C). These results raised the possibility that Apt regulates transcription of *hh*. To test the possibility, we analyzed the expression of *hh* under loss-of-function and overexpression of Apt. The expression of *hh*-lacZ and *Hh* was significantly reduced in the *apt* mutant clones in the PE (Fig. 2J-L; Fig. S3A-C) and the DP (Fig. 2M-O; Fig S3D-F). Moreover, the expression of *hh* was significantly reduced in the wing disc of larvae upon RNAi-knockdown of *apt* (Fig. S4). By contrast, overexpression of Apt increased the expression of *hh-lacZ* (Fig. 2P-R) and *hh* (Fig. S4, S5G-I). The expression of *Hh* also decreased in the *apt* mutant clones of the eye disc and the salivary gland (Fig. S5A-F). These results indicate that Apt activates the expression of *hh*.

**Fig. 2.**
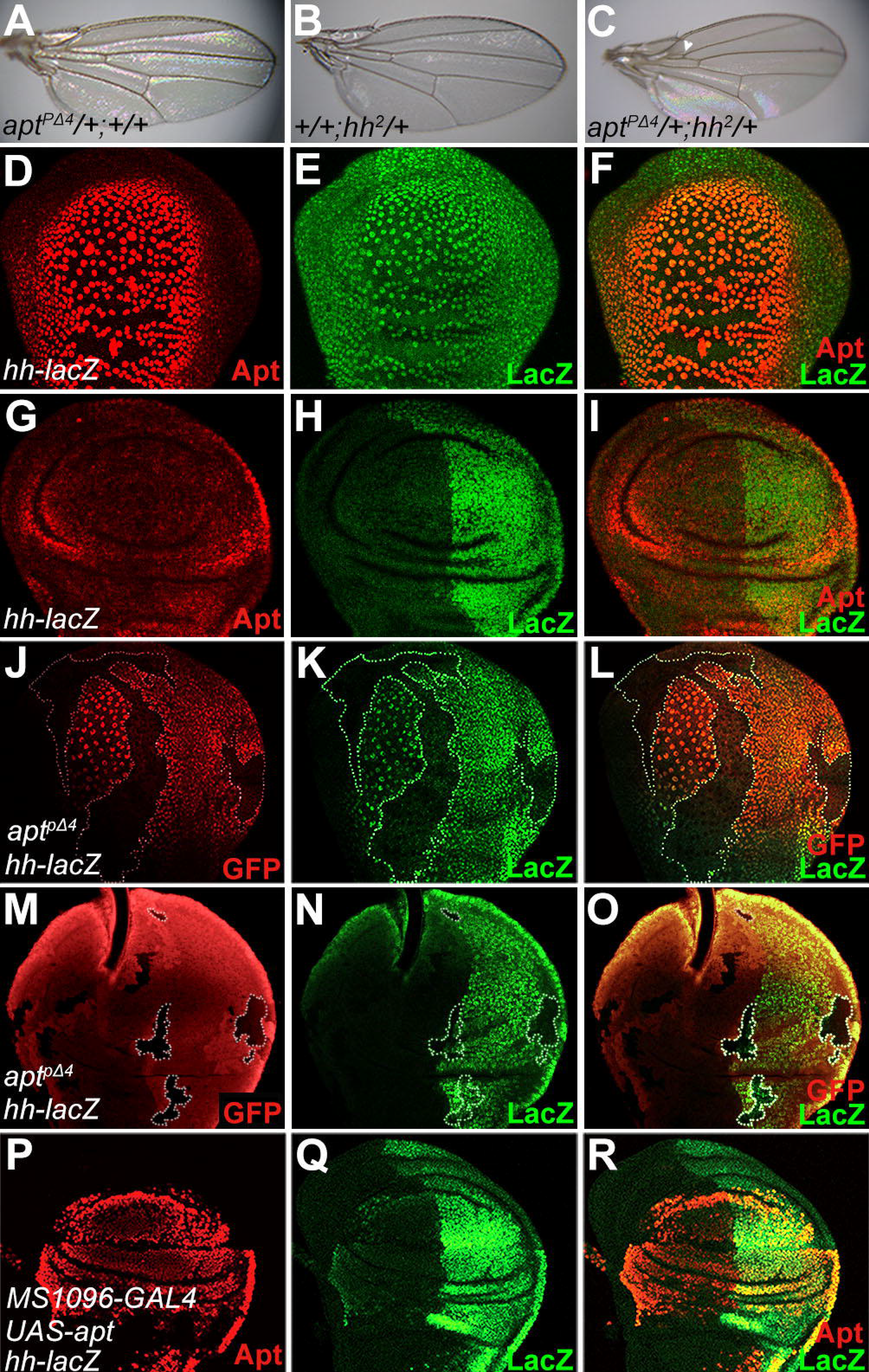
Apt regulates the expression of *hh*.

(A-C) Genetic interaction between *apt* and *hh*. Adult wings of *apt^PΔ4^/+* +/+ (A) and +/+ *hh*^2^/+ (B) show normal pattern. (C) Forty percent of *apt^PΔ4^/+ hh^2^/+* wings exhibited abnormal morphologies in anterior crossvein (ACV). An arrowhead indicates the extra ACV. Total numbers of analyzed wings were A, 158; B, 157; C, 116. (D-F) The expression of Apt (D) and *hh-lacZ* (E) in PE cells. (G-I) The expression of Apt (G) and *hh-lacZ* (H) in DP cells. (J-L) The decreased expression of *hh-lacZ* (K) in the *apt^PΔ4^* clones of the PE (J). Clones are marked by white-dotted lines. (M-O) The decreased expression of *hh-lacZ*(N) in the *apt^PΔ4^* clones of the DP (M). Clones are marked by white-dotted lines. (P-R) Overexpressed Apt (P) increased the expression of *hh* (Q).

### Apt directly controls *hh* in the wing disc

To address how Apt activates the expression of *hh*, we focused on a 15-kb region of the *hh* locus known to reproduce the normal *hh* expression pattern in the wing disc (Lee et al., 1992). We identified one potential Apt binding sequence (Liu et al., 2003) within the region (Fig. 3A). We next assessed the function of the Apt-binding site in *hh* using a CRISPR-Cas9 system (Kondo and Ueda, 2013). Since the designed gRNA contained the Apt-binding site, four Apt-binding site deletion mutants and two insertion mutants were generated (Fig. 3B-C; Fig. S6A). Homozygotes of these mutations showed reduced expression of *hh* mRNA and *Hh* protein (Fig. 3D-E; Fig. S6B) and exhibited the small wing and reduced vein 3–4 spacing phenotypes (Fig. 3F; Fig. S6C-D). Effect of *hh^ΔaptDB1^* mutation on the *hh* function was also examined under the *hh*^2^ heterozygous background. While wings of animals heterozygous for *hh*^2^ or *hh^ΔaptDB1^* were normal, transheterozygotes of *hh^ΔaptDB1^* and *hh*^2^ showed the same extra vein phenotype (Fig. 3G-I) as did transheterozygotes of *apt*-null allele and *hh*^2^. Taken together, these data suggest that Apt directly regulates the expression of *hh* in the wing disc for proper wing development.

**Fig.3.**
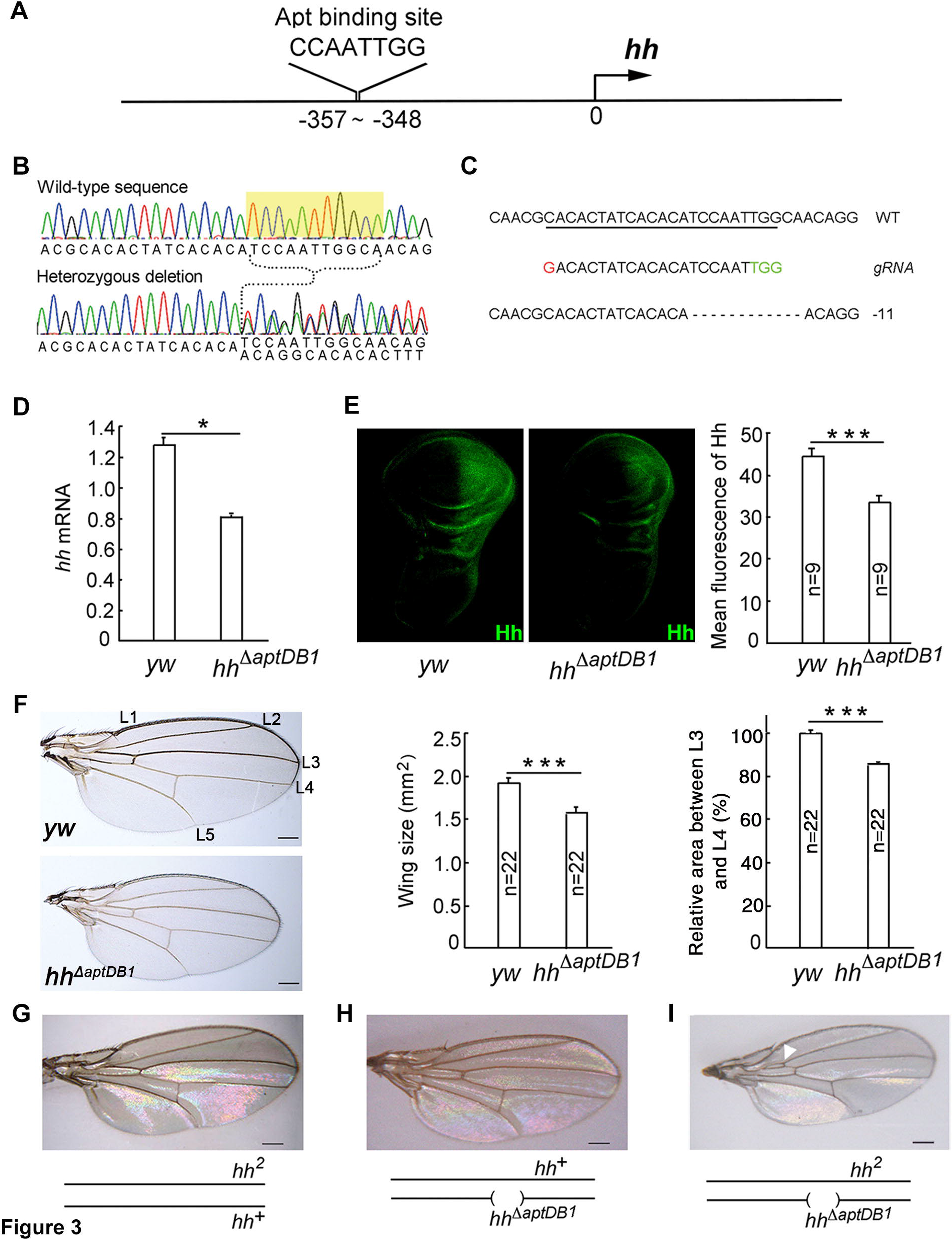
Apt directly regulates the expression of *hh* through its binding site in the *hh* promoter region.

(A) Schematic representation of the Apt-binding site in the genomic sequence of *hh*. The arrow represents transcription start site and the numbers in base pairs are distance from the start site. (B) Sequences of a wild-type allele and a heterozygous mutant of *hh^ΔaptDB1^*. The sequence of the mutant allele was inferred by subtracting a wild-type sequence from the mixed sequence. The deleted sequence is highlighted in yellow. (C) Cas9-induced mutagenesis at the *hh* locus. The *hh* locus in Cas9-induced mutants was PCR-amplified and sequenced. The wild-type sequence is shown at the top as a reference. The Cas9-gRNA target sequence is underlined with the protospacer-adjacent motifs (PAM) indicated in green. Deleted nucleotides in *hh^ΔaptDB1^* are shown as dashes. The deletion size is shown next to the sequence. (D) RT-qPCR analyses of *hh* mRNA in the wing disc of third instar larvae from *yw* or *hh^ΔaptDB1^*. Error bars, SEM from three independent experiments. Student's t tests, *p<0.05. (E) The wing disc of third instar larvae from *yw* or *hh^ΔaptDB1^* was stained with an anti-Hh antibody. The expression levels of Hh were determined by mean fluorescence. Error bars, SEM. ***p<0.001. (F-I) Deletion of the Apt-binding site in the *hh* promoter affects wing development. The wing size and the intervein region between L3 and L4 relative to that between L1 and L2 (control value was set as 100%) were decreased in *hh^ΔaptDB1^*. Error bars, SEM. Student's t tests, ***p<0.001. *hh^2^/+* (G) or *hh^ΔaptDB1^/+* (H) adult wing shows a normal phenotype. All adult wings of *hh^ΔaptDB1^/hh^2^* transheterozygotes exhibited abnormal morphologies in ACV (I). An arrowhead indicates the extra ACV. Total numbers of analyzed wings were G, 157; H, 132; I, 74. Scale bars, 200 um.

### Apt activates the *cyclin E* expression in the wing disc

We have reported that Apt induces the *cyclin E* expression in the eye disc (Liu et al., 2014).Therefore, we examined whether Apt regulates *cyclin E* also in the wing disc. To do this, we first performed a double-staining experiment. In the wild-type wing disc, Apt and *cyclin E* were co-expressed (Fig. 4A-C). Furthermore, the expression of *cyclin E* was significantly reduced in the *apt* mutant clones (Fig. 4D-F; Fig. S7A). The expression of *cyclin E* mRNA was also reduced upon RNAi-knockdown of *apt* in the wing disc (Fig. S7B). By contrast, the expression of *cyclin E* and its mRNA was increased by overexpression of Apt in the wing disc using *MS1096-Gal4* and *UAS-apt* (Fig. 4G-I; Fig. S7B). These results indicate that Apt activates the expression of *cyclin E*in the wing disc.

**Fig. 4.**
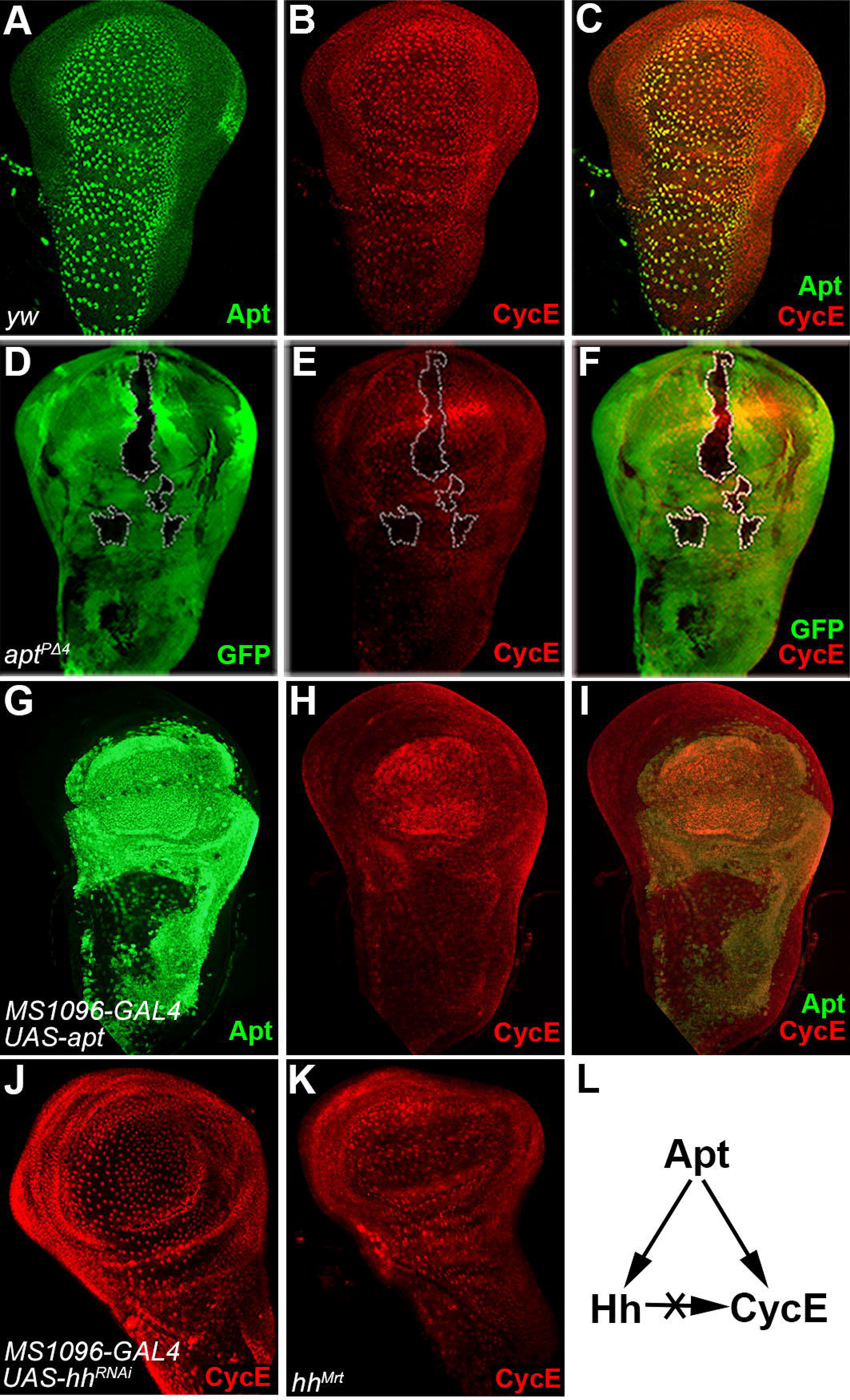
Apt controls the expression of *cyclin E*.

(A-C) The expression of Apt (A) and *cyclin E* (B) in the PE of control wing disc. (D-F) Decreased *cyclin E* (E) expression in the *apt* mutant clones (D). (G-I) Overexpressed Apt with MS1096-GAL4 (G) increased the expression of *cyclin E* (H). (J, K) The expression of *cyclin E* in the wing disc from the *hh^RNAi^* knockdown (J) and *hh* gain of function mutant animals (K). (L) Regulation of *hh* and *cyclin E*by Apt in the wing disc. Wing discs from third instar larvae of wild-type (A-C), *hs-FLP; FRT42D, Ubi-GFP/FRT42D, apt^PΔ4^* (D-F), *MS1096-GAL4; UAS-apt* (G-I), *MS1096-GAL4; UAS-hh^RNAi^* (J) and *hh^Mrt^* (K) animals were stained with the anti-Apt antibody (A, G), the anti-GFP antibody (D), the anti-cyclin E antibody (B, E, H, J and K). (C, F, I) Merged images of A and B, D and E, G and H, respectively.

We then asked whether the regulation of *cyclin E* by Apt is mediated through *Hh*. To test this idea, we compared the expression of *hh* and *cyclin E* upon overexpression of Apt in the wing disc. Both *hh* and *cyclin E* were induced by overexpression of Apt (Fig. 2Q,4H). However, their expression patterns were different. *cyclin E* was induced in all region of the wing disc, whereas the expression of *hh* was restricted in the posterior compartment. Moreover, the expression of *cyclin E* was not changed by RNAi-knockdown of *hh* using *MS1096-Gal4* and *UAS-hh^RNAi^* (Fig. 4J) and in an *hh* gain of function mutant *hh^Mrt^* that exhibits ectopic expression of *hh* in the anterior compartment (Tabata and Kornberg, 1994) (Fig. 4K). These data suggest that the activation of *cyclin E* by Apt is independent of *Hh* in the wing disc (Fig. 4L).

### Apt directly controls *cyclin E* in the wing disc

Since Apt directly activates the expression of *cyclin E* in the eye disc (Liu et al., 2014), we anticipated a direct role of Apt in the expression of *cyclin E* also in the wing disc. This expectation was verified by transgenic reporter assays. The reporter gene (Liu et al., 2014) carries the endogenous promoter and the *cyclin E* regulatory element containing a wild-type Apt-binding site (*cycEPlacZ*) (Fig. 5A) or a mutated site (*cycEMPlacZ*) (Fig. 5E). Although *cycEPlacZ* with the wild type binding site recapitulated the *cyclin E* expression in the wing disc (Fig. 5B-D), base substitutions in the Apt-binding site in *cycEMPlacZ* abolished the lacZ expression (Fig. 5F-H). These results indicate that Apt directly activates *cyclin E* through its binding site in the regulatory region of *cyclin E.*

**Fig. 5.**
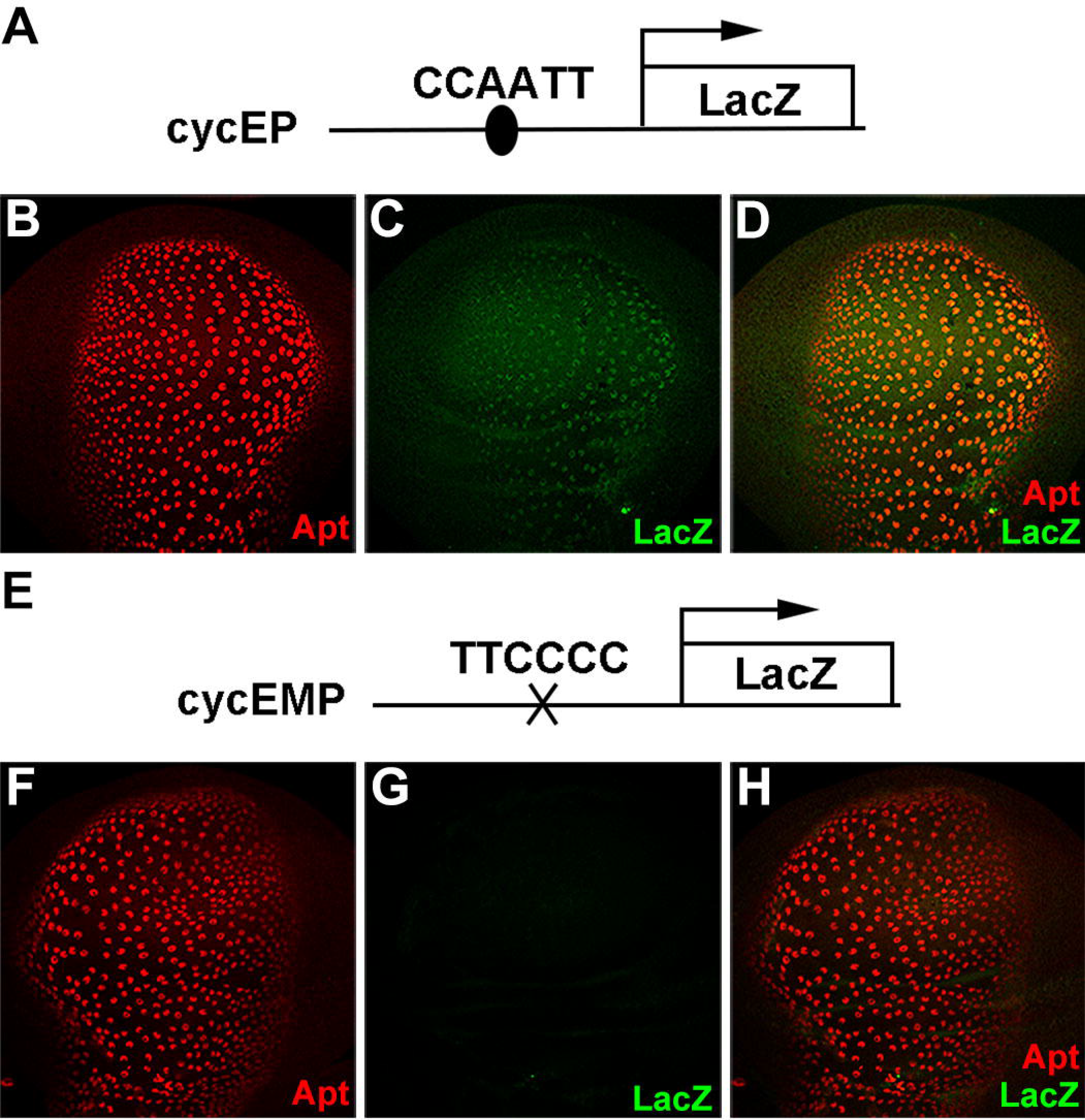
Apt directly regulates the expression of *cyclin E* through its binding site in the *cyclin E* promoter region.

(A) Schematic illustration of the lacZ reporter construct driven by the *cyclin E* promoter carrying the wild type Apt binding site. (B-D) The reporter *cycEPlacZ* (C) was coexpressed with the endogenous Apt (B) in the wing disc. (E) Schematic illustration of the lacZ reporter construct driven by the *cyclin E* promoter carrying the mutant Apt binding site. (F-H) The expression of reporter *cycEMPlacZ* (G).Wing discs of third instar larvae were stained with anti-Apt antibody (B and F) and anti-β-galactosidase antibody (C and G). (D and H) Merged images of B and C and of F and G, respectively.

### Apt controls cell proliferation by inducing *hh* and *cyclin E*

Because both *Hh* and *cyclin E* are involved in cell proliferation (Jiang and Hui, 2008;Knoblich et al., 1994), defects in *apt* would affect the cell number in the wing disc. As expected, we observed significant decrease in the cell number in an *apt* mutant clone using DAPI staining (Fig. 6B). Moreover, phalloidin labeling revealed disruption of the linear arrangement of cells in the clone (Fig. 6C). When Apt was overexpressed in the wing disc, the number of DAPI-stained cells was not significantly changed from that in the control disc (compare Fig. 6F with 6D). However, the linear arrangement of cells was disrupted (compare Fig. 6G with 6E). Since *Hh* and *cyclin E* are required for the regulation of apoptosis (Guerrero and Ruiz i Altaba, 2003;Hwang and Clurman, 2005;Ruiz i Altaba, 1999), we asked whether the overexpression phenotypes are caused by apoptosis. To test this, we investigated apoptosis in wing discs by staining with anti-Caspase-3 antibody. In the third instar wing disc from *apt* mutant clones and wild type, few apoptotic cells were observed (Fig. 6H-J). However, in the wing disc from an Apt-overexpressed larva, the number of apoptotic cells was significantly increased (Fig. 6K). This presumably explains why wing size was reduced upon overexpression of Apt (Fig. 1G).

**Fig. 6.**
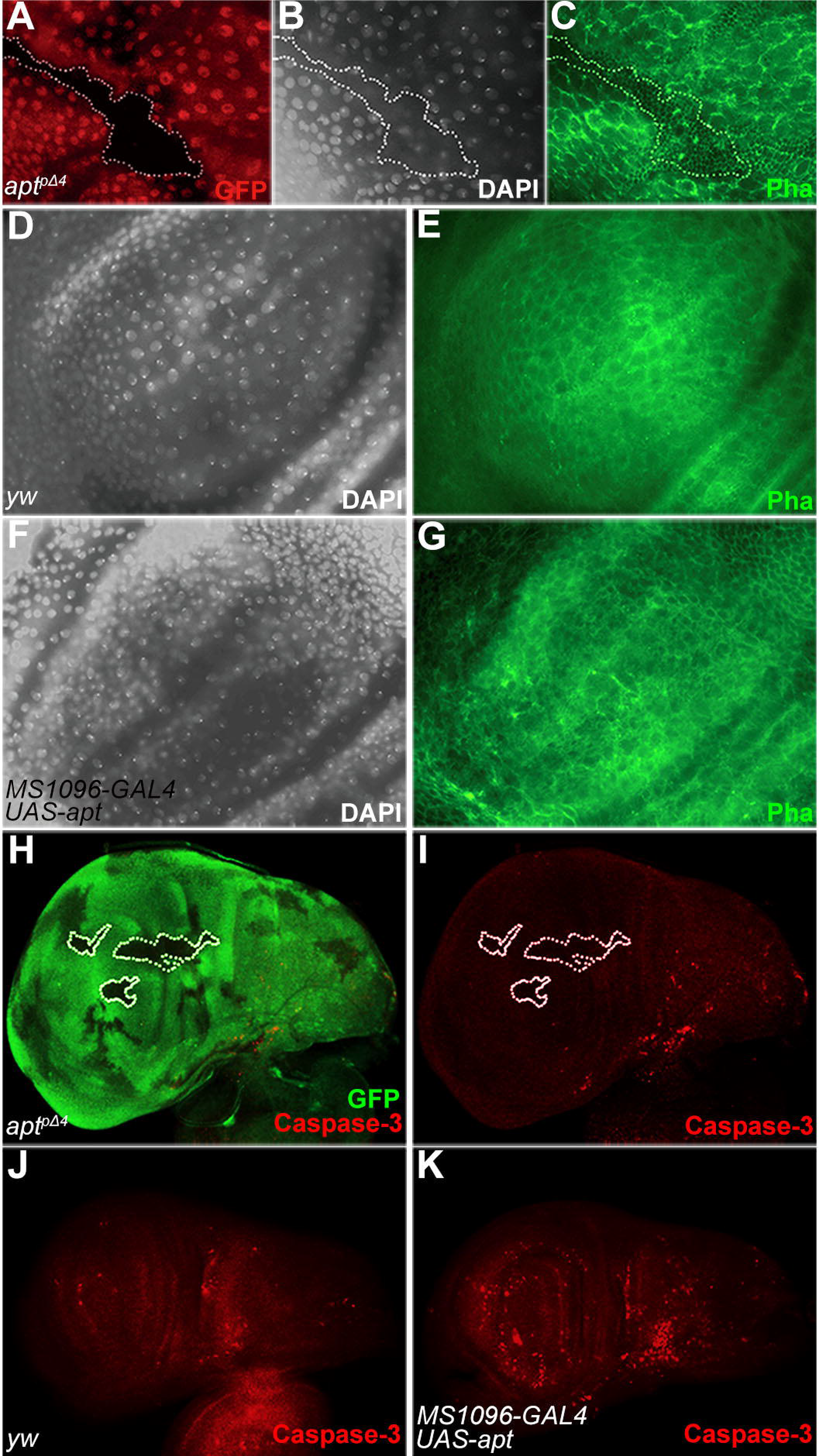
Apt is required for production of proper cell number and arrangement of wing discs.

(A-C) Cell number (B, marked by DAPI and array of cells (C, marked with phalloidin) were affected in an *apt* mutant clone (A, lack of the GFP signals and marked with a broken line. (D, E) Cell number (D) and array (E) from a wild-type wing disc were visualized with DAPI (D) and phalloidin (E). (F, G) Overexpression of Apt in the wing pouch resulted in slight decrease in the cell number (F) and irregular arrangement of cells (G). (H-J) Apoptosis was barely detectable in the *apt* mutant clones (H, I) and wild-type wing disc (J). (K) Overexpressed Apt in the wing disc increased the number of apoptotic cells. The wing discs of *hs-FLP; FRT42D, Ubi-GFP/FRT42D, apt^PΔ4^* (A-C, H, I), wild-type (D, E and J) and *MS1096-GAL4; UAS-apt* (F, G and K) animals were stained with the anti-GFP antibody (A and H), DAPI (B, D and F), phalloidin (C, E and G) and Caspase-3 antibody (H-K).

Homozygotes of *hh* mutations for the Apt-binding site exhibited the small wing but not the blistered phenotype. However, *hh* and *cyclin E* double mutant recapitulates the smaller and blistered wing. While *CycE^2^/+* flies showed normal wings, three percent of *hh^bar3^*/*hh^bar3^* and eighteen percent of *CycE^2^/+ hh^bar3^/hh^bar3^* flies showed the smaller and blistered phenotypes (Fig. S8A-C). We also observed genetic interaction between *hh* and *cyclin E* in the extra crossvein phenotype. While *CycE^JP^*/+ and *hh^2^*/+ flies showed normal wings, fifty-four percent of *CycE^JP^*/+ *hh^2^*/+ flies exhibited wings with the extra crossvein (Fig. S8D-F). Collectively, these data suggest that Apt controls wing development by inducing appropriate amounts of *Hh* and *cyclin E.*

## DISCUSSION

Here, we revealed that the transcription factor Apt regulates *Drosophila* wing development, at least in part, through directly activating the expression of *hh* and *cyclin E* to control wing patterning and growth. Both loss-of-function and gain-of function assays clearly demonstrated that Apt is vital for wing development. Further studies showed that knockdown of *apt* attenuated, while overexpression of *apt* activated the expression of *hh* and *cyclin E.*

In the wing disc, *Hh* exclusively expresses in the P compartment. After many modifications, the mature *Hh* ligands are secreted from the P compartment and reach ~12cell rows near A/P boundary of the A compartment (Basler and Struhl, 1994; Tabata et al., 1992;Tabata and Kornberg, 1994). Ci expresses solely in the A compartment (Slusarski et al., 1995). Without the *Hh*, full-length Ci is ubiquitinated by SCF^Slimb^ to partial degradation, culminating in formation a truncated transcriptional repressor termed Ci^R^. Ci^R^ enters into the nucleus to repress the expression of *hh* in the A compartment (Aza-Blanc et al., 1997;Jiang and Struhl, 1998;Smelkinson and Kalderon, 2006). In this study, we found the ubiquitous expression of Apt in the wing disc (Fig. 1A,B). However, the expression of *hh* is restricted in the P compartment of DP cells. Overexpression of *apt* in the wing disc with the MS1096-GAL4 driver emerges the ectopic expression of *hh* in the A compartment, suggesting that Apt is sufficient to turn on *hh* expression. We speculate that during the normal development progress, Apt might cooperates with others factors (such as Ci^R^) to restrict the expression region of *hh*. It is interesting to investigate the relationship between Ci^R^ and Apt.

To assess the importance of the Apt-binding site in the promoter region of *hh*, we first tried a transgenic reporter assay. However, the regulatory region of *hh* encompassing the upstream region and the 1st intron (~15 kb) (Lee et al., 1992) is too large to make a reporter construct for conventional P-element mediated transgenesis. Therefore, we employed the CRISPR-Cas9 system (Kondo and Ueda, 2013) to mutagenize the endogenous Apt-binding site in the *hh* promoter. All 6 independent mutants exhibited the same phenotypes (reduced expression of *hh*, reduced wing size and the space between L3 and L4), suggesting that the observed phenotypes are not due to off-target effect of Cas9. Nevertheless, we inspected the possibility of off-target effect. Since our gRNA carries the binding sequence for Apt, a binding site of Apt in other than the *hh* promoter could be the most likely candidate for off-target. However, all the 6 mutants showed the wild type sequence around the Apt-binding site in the *cyclin E* promoter (Fig. S9). Furthermore, transheterozygotes between *hh^2^* and *hh^ΔaptDB1^* exhibited the *hh* mutant phenotype, smaller wing with extra crossvein. Taken together, these data strongly suggest that the observed phenotypes are not due to off-target effect.

Although our data strongly support that Apt is a transcription factor of *hh*, mutating the Apt binding site on *hh* promoter does not induce severe phenotypes. Beside Apt, other factors might also regulate *hh* transcription. Therefore, both knockdown and overexpression of *apt* only moderately affect the expression of *hh*. *Hh*, an important morphogen, plays multifaceted roles in segmentation and wing patterning. Previous findings paid more attention on the protein modification of *Hh*. The mechanism underlying *hh* transcription is not clear. Here our studies unveil that Apt acts as a transcription factor of *hh*.

While *Hh* has been implicated in induction of *cyclin E* through Ci (Duman-Scheel et al., 2002), subsequent researches have shown that *cyclin E* accumulates in the *Mad^1–2^Su(H)ci* mutant cells (Firth and Baker, 2005). So whether *Hh* activates *cyclin E*is controversial. In this study we showed that RNAi-mediated knockdown of *hh* or ectopic expression of *hh* in the anterior compartment did not change the expression of *cyclin E.* Taken together, these observations argue against the notion that *hh* regulates *cyclin E*in the wing disc.

Hyperactivation of *Hh* pathway has been complicated in many tumors (Clement et al., 2007;Jiang and Hui, 2008). It will be fruitful to investigate whether Apt is upregulated in *Hh*-related tumors. The previous work indicates that Apt involves in tumorigenesis (Woodhouse et al., 2003). It is also interesting to explore whether Apt regulates tumorigenesis through activating *Hh* signaling. Our finding that Apt regulates wing development through activating *hh* raises a possibility that Apt acts as a potential clinical target for *Hh*-related tumors.

## MATERIALS AND METHODS

### *Drosophila* Strains

Strains used were as follows. *apt^pΔ4^* (Eulenberg and Schuh, 1997), *apt^p2^* (Liu et al., 2003), *cycEPlacZ*and *cycEMPlacZ*(Liu et al., 2014), *UAS-apt* (gift of D. Montell), *UAS-GFP* (gift of Y. Hiromi). *hh^Mir^* was obtained from *Drosophila* Genetic Resource Center. *hh^2^*, *hh^Mrt^*, *hh^bar3^*, *CycE^2^*, *CycEf^P^*, *hh-LacZ, MS1096-GAL4, sd-GAL4, dpp-Gal4, ptc-GAL4* and *UAS-hh^RNAi^* were obtained from Bloomington *Drosophila* Stock Center. *UAS-apt^RNAi^* was obtained from Tsinghua Fly Center. *y^2^cho^2^v^1^; attP40{nos-Cas9/CyO, y^1^ v^1^ P{nos-phiC31\int.NLS}X; attP40, y^2^ cho^2^ v^1^, y^2^ cho^2^ v^1^/Y^hs-hid^; Sp/CyO, y^2^ cho^2^ v^1^; PrDr/TM6C* were obtained from NIG-Fly.

### Clonal analysis

Homozygous *apt* loss-of-function clones were generated by *hs-FLP/FRT* recombination (Theodosiou and Xu, 1998). *FRT42D* and *apf^A4^/CyO* were recombined to generate *FRT42D, apf^A4^.* Six pairs of *FRT42D, apf^A4^* cross to *Gla/CyO* were allowed to lay eggs in G418-containing medium, and then test each line with *apt^P2^/CyO. hs-FLP; FRT42D, Ubi-GFP/CyO* crossed with *FRT42D, apt^PA4^/CyO* were performed at 25°C. Heat shocks were performed 32–56 hours after egg-laying for 1.5 hours at 37.5°C.

### Generation of CRISPR constructs

To induce mutations in the Apt-binding site in the *hh* promoter region, we used a Cas9-gRNA system. We designed gRNA in the *hh* promoter region carrying the binding sequence of Apt (Fig. 3A). The corresponding sequence was introduced into the pBFv-U6.2 vector and the gRNA transgenic flies were generated as described (Kondo and Ueda, 2013). gRNA females were crossed to Cas9 males to obtain the founder animals. Male founders were crossed to female balancer. Offspring male flies were balanced and stocked. Genomic DNA was extracted from each offspring male and used for molecular characterization. PCR primers were designed to construct gRNA expression vectors and to amplify the promoter region of *cyclin E* (Table S1).

### RT-qPCR analysis

Wing discs were dissected from 40 third instar larvae. Total RNA was prepared from the dissected tissues using an RNAprep Pure Tissue kit TIANGEN #DP431). cDNAs were synthesized using a Prime Script^TM^||1^st^ strand cDNA synthesis kit (TaKaRa #6210A). qPCR was conducted with Bio-Rad CFX96 real-time system using a SuperReal PreMix Plus (SYBR Green) Kit (TIANGEN #FP205) in a 20 ul reaction containing 2 pmol of relevant primers. The amount of mRNA was normalized to that of control tubulin mRNA. PCR primers were designed to amplify the *hh* region (Table S1).

### Antibodies and Immunohistochemistry

Staining of larval tissues was performed as described previously (Liu et al., 2014). Larvae were dissected in PBS, fixed in 25 mM PIPES-KOH (pH 7.0), 0.5 mM EDTA, 0.25 mM MgSO4 and 4% formaldehyde for 40 minutes on ice and then permeabilized for 15 minutes at room temperature in PBS containing 0.5% NP-40. The following primary antibodies were used in overnight incubations at 4°C in blocking solution: rabbit anti-Apt (1:1000) (Liu et al., 2014), rabbit anti-Hh (1:800, gift of T. Tabata), rabbit anti-GFP (1:200, Molecular Probes), mouse anti-GFP (1: 400, Molecular Probes), rabbit anti-β-galactosidase (1:2000, Cappel), rabbit Caspase3 (1:50, Cell Signaling Technology), mouse anti-β-galactosidase (1:500, Sigma), FITC-conjugated phalloidin (1:200, Sigma), mouse anti-Ubx (1:10, Developmental Studies Hybridoma Bank (DSHB)), goat anti-cyclin E (1:200, Santa Cruz). The secondary antibodies used were as follows: Alexa 488 donkey anti-rabbit IgG conjugate (1:500, Molecular Probes), Alexa 488 donkey anti-mouse IgG (1:500, Molecular Probes), Cy3-conjugated donkey anti-mouse IgG (1:500, Sigma), Cy3-conjugated goat anti-rabbit IgG (1:500, CWBIO), bovine anti-goat IgG-CFL555 (1:500, Santa Cruz). Mounting used VECTASHIELD Mounting Medium with DAPI (Vector Labs). The caspase-3 staining was did as described previously (Rudrapatna et al., 2013).

### Microscopy and Image Treatment

Images were acquired in Leica TCS SP5 confocal microscope and Olympus cellSens, treated with Adobe Photoshop CS6 image programs. Wing size and space between vein 3 and vein 4 or that between vein 1 and vein 2 were measured on each picture using the ImageJ computer program.

### Statistical analysis

Results are given as means SEM; each experiment included at least three independent samples and was repeated at least three times. Group comparisons were made by two-tailed unpaired Student's t-tests. *P < 0.05; **P < 0.01, and ***P < 0.001.

## Acknowledgements

We thank Denise J. Montell, Tetsuya Tabata, Jiong Chen, Yasushi Hiromi, Shigeo Hayashi, Ryu Ueda, Shu Kondo and Hua Tang, Tsinghua Fly Center, NIG-Fly, Kyoto stock and Bloomington Stock Center for providing antibodies, fly strains and technical advice.

## Author Contributions

X.F.W., S.H., and Q.X.L. designed research, X.F.W., Q.C., and C.L.F. performed experimlents and X.F.W., Z.Z.Z., S.H., and Q.X.L. analyzed data and wrote the manuscript.

## Supplementary information

Supplementary information available online at dev.biologists.org/lookup/suppl

